# A novel high-throughput molecular counting method with single base-pair resolution enables accurate single-gene NIPT

**DOI:** 10.1101/597732

**Authors:** David S. Tsao, Sukrit Silas, Brian P. Landry, Nelda Itzep, Amy B. Nguyen, Celeste K. Kanne, Vivien A. Sheehan, Rani Sharma, Rahul Shukla, Prem N. Arora, Oguzhan Atay

## Abstract

Next-generation DNA sequencing is currently limited by an inability to count the number of input DNA molecules. Molecular counting is particularly needed when accurate quantification is required for diagnostic purposes, such as in single-gene non-invasive prenatal testing (sgNIPT) and liquid biopsy. We developed Quantitative Counting Template (QCT) molecular counting for reconstructing the number of input DNA molecules using sequencing data. We then used QCT molecular counting to develop sgNIPT of sickle cell disease, cystic fibrosis, spinal muscular atrophy, alpha-thalassemia, and beta-thalassemia. Incorporating molecular count information into a statistical model of disease likelihood led to analytical sensitivity and specificity of >98% and >99%, respectively. Validation of sgNIPT was further performed with maternal blood samples collected during pregnancy, and sgNIPT was 100% concordant with newborn follow-up.

## Introduction

Next-generation sequencing (NGS) has revolutionized both research and clinical practice. In 2001, a global effort was required to sequence the first human genome; nowadays over 100,000 human genomes can be sequenced in a single center.^1^ In many biological subfields, significant advances have been made by innovative experimental designs for generating data resolvable by NGS.^2, 3^ In the clinic, performing whole exome sequencing is now considered the standard of care for any undiagnosed congenital and neurodevelopmental disorders.^4^

Translation of genome sequencing technology to the clinic has been even more widely adopted in prenatal and oncology care. Now that targeted oncologic therapies are indicated only when specific genetic profiles are found in the tumor, DNA sequencing of biopsied tissue is FDA-approved and covered by national insurance programs.^5, 6^ More recently, it has been discovered that circulating tumor DNA (ctDNA) can be found in the cell-free DNA (cfDNA) purified from plasma.^7^ This has led to “liquid biopsies” that detect cancer mutations by DNA sequencing of circulating cell-free DNA (cfDNA) purified from a blood sample.^8^ In prenatal care, cfDNA of fetal origin is used to detect fetal aneuploidies as early as the 10th week of gestation from maternal blood.^9, 10^ These non-invasive prenatal tests (NIPT) are routinely used in clinical care, and they are offered to all pregnancies via national health insurance.^11^

However, these cfDNA-based tests have been limited to detection of chromosomal abnormalities or large structural variants in prenatal testing^10^ and late-stage cancers in liquid biopsy.^12^ Yet, several prevalent genetic disorders (e.g. hemoglobinopathies, cystic fibrosis, and spinal muscular atrophy) are caused by single nucleotide variants (SNV) or a single gene copy number change. These recessively inherited disorders are particularly challenging to detect because they require precise quantification of fetal cfDNA over a significant background of identical maternal cfDNA. In liquid biopsy, even though copy number variation is a hallmark of cancer,^13^ current FDA-approved diagnostics are limited to resolving a 30% to 120% increase in the copy number found in cfDNA and therefore are only applicable to late-stage cancers.^14–16^ These limitations are largely due to traditional library preparation methods for NGS that introduce significant levels of noise that obscure the relationship between DNA molecular counts of the input sample and the final sequencing read count. Without the ability to perform absolute quantification, NGS cannot reliably quantify fetal SNVs in prenatal testing, detect gene amplifications that are present in Stage 1-2 cancers, or sensitively detect rare cell-free tumor DNA sequences that may only be present at 1-10 molecules in the sample.^17^ Importantly, increased read depth does not increase sensitivity in these contexts. Amplification and/or enrichment biases can result in 10-fold variability in output read-counts across loci, while accurate determination of single-gene NIPT or early stage liquid biopsy may require quantification of input DNA molecular counts at differences <5%. In the vast majority of cases, the major sources of noise are introduced during amplification and library preparation, and therefore increased sequencing depth does not result in improved accuracy.^18^

Currently, the only method to obtain accurate absolute quantification of DNA molecules is via digital PCR (dPCR).^19, 20^ While dPCR has been a revolutionary technology, especially for developing reference materials, its limited multiplexability of 2-4 variants at a time significantly limits its clinical use. Moreover, since dPCR is not sequencing-based, it is unable to resolve between multiple, adjacent variants as is commonly the case in single gene disorders and oncogenes.

We developed a sequencing-based molecular counter with single base-pair resolution and vastly superior multiplexing ability. Since molecular abundance information in sequencing read depth data is typically corrupted by library preparation, we developed a technique to encode molecular abundances prior to PCR amplification and subsequent library preparation steps using Quantitative Counting Template (QCT) molecules. Absolute quantification of DNA molecules is then decoded via customized bioinformatic analyses. We applied our molecular counter technology to single-gene NIPT for hemoglobinopathies, cystic fibrosis, and spinal muscular atrophy. These recessively inherited disorders require high-accuracy molecular count information to be able to determine whether a particular mutation is elevated relative to the maternal background.

Medical guidelines recommend that all pregnancies should be screened for these disorders.^21, 22^ Hemoglobinopathies, i.e., sickle cell disease, beta-thalassemia, and alpha-thalssemia, are the most common genetic disorders in the world, affecting more than 300,000 births each year.^23^ The number of pregnancies that are at-risk is even higher, with 7% of pregnant women globally carrying a significant pathogenic variant for hemoglobinopathies alone.^23^ In the US, 7% of African-Americans^24^ and 1.5% of all newborns are carriers for sickle cell disease (sickle cell trait, SCT).^25^ The carrier rates for cystic fibrosis and spinal muscular atrophy are 3% and 2%, respectively.^26, 27^ Although NIPT for chromosomal abnormalities is widely used, NIPT for recessively inherited single gene disorders are not currently available. Rather, the only way to diagnose these disorders in the fetus is through invasive methods such as amniocentesis or CVS, which carry significant miscarriage risk.^28^

In this study we showed that our QCT method accurately counts molecules with single base-pair resolution. We then applied QCT molecular counting to NIPT for mendelian disorders, including sickle cell disease, thalassemias, cystic fibrosis, and spinal muscular atrophy. These NIPTs require only a single 10mL sample of maternal blood (Fig. 1). We validated our NIPT with preclinical samples (estimated sensitivity of >98%+ and specificity of >99%) comprised of genomic DNA sheared to mimic the fragmentation of cfDNA. Crucially, reliable NIPT results were only obtainable via accurate molecular counting of these genes. NIPT assays were also performed on non-pregnant cell-free DNA control samples to show the concordance of sheared DNA samples with cfDNA. Finally, we performed NIPT on maternal blood samples from 37 pregnancies. All NIPT results were 100% concordant with the newborn genotype, even in a challenging sample with fetal fraction as low as 5%.

**Figure 1.**
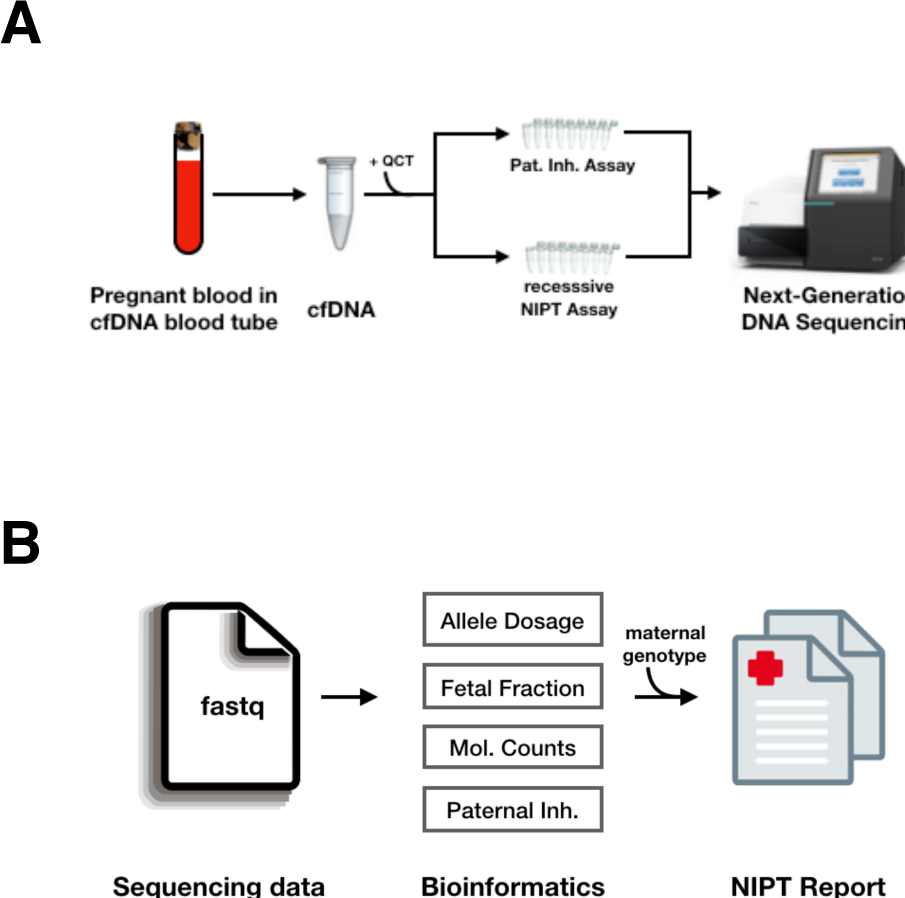
Overview of single-gene NIPT. (A) Clinical workflow uses a single blood tube of maternal blood. Amplicon-based assays are performed and sequenced using an Illumina Miseq instrument. (B) Bioinformatic analyses recover the dosage of pathogenic alleles, the fraction of cfDNA isolated from maternal blood that is of fetal origin, the number of DNA molecules assayed, and paternal inheritance of variants not found in the mother’s genotype. These analyses are combined with the maternal genotype to perform a statistical analysis of fetal genotype, resulting in an NIPT report.

## Results

### High throughput molecular counting with single base pair resolution

We first developed a technique to count the number of DNA molecules in a PCR using amplicon NGS workflows (Fig. 2). In this assay, a number of Quantitative Counting Template (QCT) synthesized DNA molecules are spiked-in to the cell-free DNA specimen prior to PCR amplification. The QCT sequence is designed to co-amplify at the same rate with its corresponding gene of interest by incorporating homology regions, especially in PCR priming sites (Fig. 2A). QCTs are designed such that the number of molecules added can be independently calculated from sequencing data. The relationship between read depth and molecular counts is then used to also determine gene of interest molecular count in the input sample (Fig. 2B).

**Figure 2.**
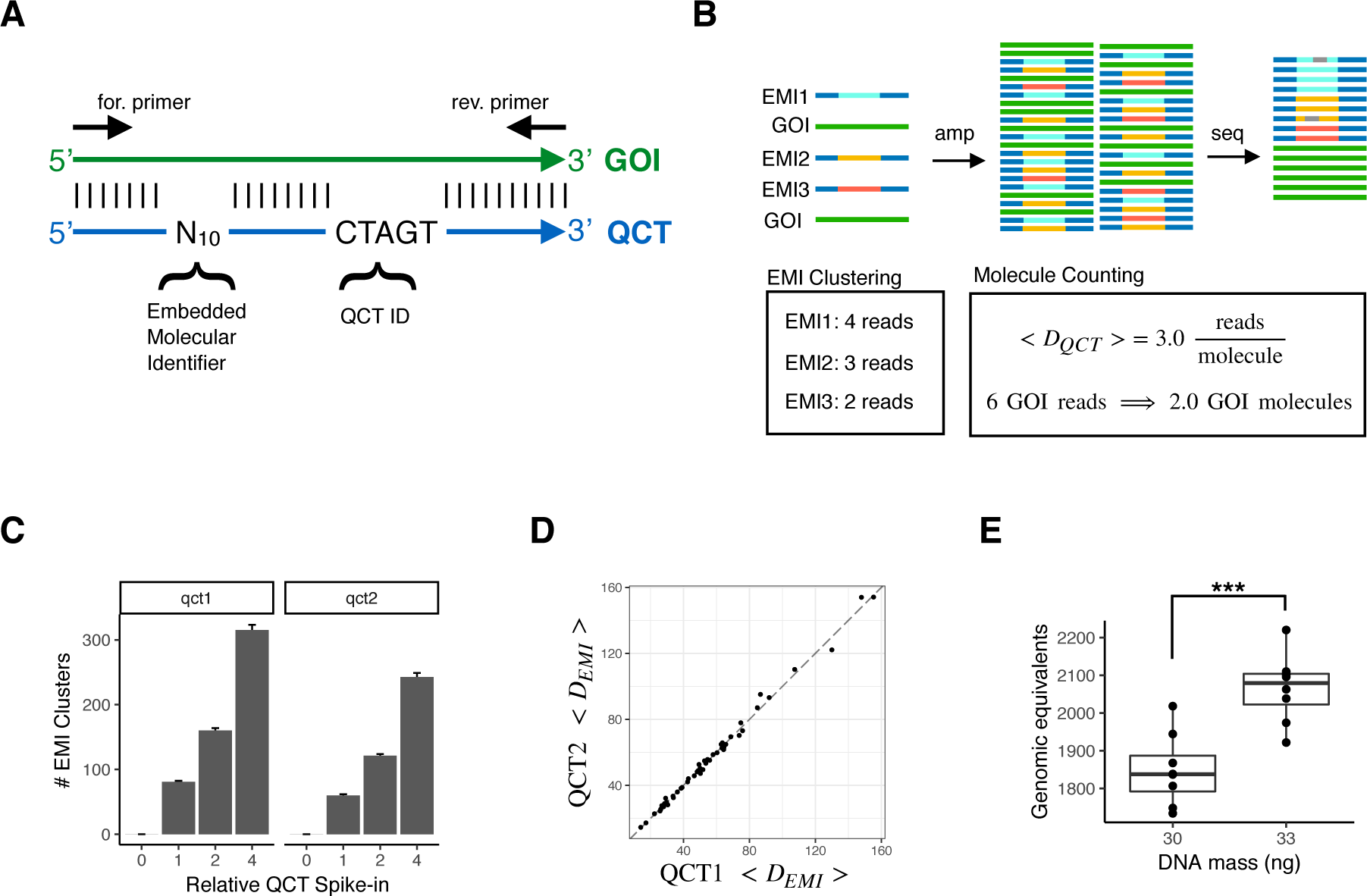
Accurate measurements of assayed genomic equivalents by Quantitative Counting Template (QCT) molecules. (A) Sequence design of QCT molecule pools. QCTs are designed to co-amplify with a gene of interest (GOI). An Embedded Molecular Identifier comprised of randomized bases ensures each QCT molecule has a unique sequence. (B) Schematic of QCT molecule counting analysis. QCT molecules are added to the DNA sample, and the mixture is amplified and sequenced. EMI reads may have sequencing errors (gray bars), which are identified and corrected for by clustering analysis. The number of GOI molecules in the sample is calculated from the GOI read depth and the average number of reads per EMI cluster. (C) Pools QCT1 and QCT2 were synthesized with different QCT IDs and diluted to approximately 100 molecules at 1x dilution. The barplot shows the mean across 24 PCR replicates at each dilution factor. Error bar is 1 standard error of the mean. (D) The average depth per QCT1 molecule and QCT2 molecule in a PCR tube is shown for the 48 PCR replicates at dilution factors 1x and 2x. Dashed line has slope=1, intercept=0. (E) 30ng or 33ng of sheared genomic DNA was added to >100 QCTs, amplified, and sequenced. At 30ng, the assayable sheared DNA mean was 1849 molecules (CV = 5.2%, n=8); at 33ng the mean was 2066 molecules (CV=4.4%, n=8). The mean GE measured for 33ng was significantly greater that that for 30ng (*p* = 0:00019, one sided t-test).

To facilitate counting of the number of QCT molecules spiked-in to each tube, the QCT sequence contains a barcode comprised of 10 randomized bases in which A, C, T, or G is stochastically incorporated during oligo synthesis (Fig. 2A). Because up to 4^10^ QCT sequences are synthesized in a pool and only ∼100 to 1000 QCT molecules are spiked-in to a PCR amplification, it is exceedingly unlikely for any two QCT molecules to have the same sequence. The randomized sequence of each molecule thus comprises an Embedded Molecular Index (EMI) that identifies the molecule and its amplification progeny (Fig. 2B). After PCR amplification and DNA sequencing, the number of QCT molecules added to each specimen should correspond to the number of EMI sequence clusters observed, and the gene of interest molecular count is determined.

To test our ability to count the number of QCT molecules present in a reaction, we added 4x, 2x, 1x, or 0x QCT molecules to PCRs, with 1x corresponding to approximately 100 QCT molecules. Twenty-four replicates were assayed at each spike-in level, for a total of 96 reactions. Embedded molecular indexes from the same PCR tube that differ by up to 2 mismatches were clustered together in order to avoid counting spurious indexes that arise due to sequencing error. We observed the expected relationship in index clusters detected by sequencing and the 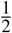 dilution series of QCT molecules (Fig. 2C). The number of QCT molecules added at each level is expected to vary among replicates due to sampling noise; addition of *N* molecules to a reaction should exhibit a standard deviation 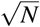 according to Poisson statistics. This effect was observed in our data, as the variance of QCT molecule counts increased with more molecules added (Fig. 2C).

We confirmed that the EMI cluster numbers were robust to read-depth variation by repeating cluster analysis on sequencing data that had been subsampled to 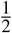 of its original read depth (Fig. S1). The number of EMI clusters obtained from subsampled data matched nearly perfectly with the full sequencing data in the vast majority of cases. In several PCR assays, subsampling the reads resulted in an underestimate of EMI clusters compared to the full read data. This effect did not occur in assays with >10 reads per EMI cluster on average, suggesting that a sequencing depth of at least 10 reads per molecule is sufficient for robust results.

Quantification of assayable human genomic equivalents (number of haploid genome copies in the sample) relies on the assumption that the gene of interest and QCT DNA co-amplify at exactly the same rate. Therefore, QCT sequences are designed to use the same PCR primer binding sites and generate the same length amplicon as the gene. However, since the internal sequence differs between the QCT and gene, QCT molecules could hypothetically be amplified at a different rate compared to the gene. To ensure that the PCR amplification is robust to sequence composition, we examined the average depth per molecule, 〈*D_QCT_*〉, of two QCT pools. Pools QCT1 and QCT2 were synthesized with QCT IDs TCGCC and CTAGT, respectively. Approximately 100-200 molecules of each QCT pool were added per PCR, and the average read depth per molecule from each pool was compared across 48 PCRs (Fig. 2D). Since QCT pools incorporate 10 randomized bases in addition to the 5 mismatches in the QCT IDs, the sequences of the ∼100 QCT molecules from each pool in a PCR assay are highly diverse. In each PCR assay, 〈*D_QCT_*_1_〉 and 〈*D_QCT_*_2_〉 were highly consistent. 〈*D_QCT_*_1_〉 spanned an 8-fold range across PCR assays due to endpoint PCR variance and uneven normalization prior to sequencing. Nonetheless, 〈*D_QCT_*_2_〉 was highly correlated with 〈*D_QCT_*_1_〉 with *R*^2^ > 0.99, and the mean difference between 〈*D_QCT_*_1_〉 and 〈*D_QCT_*_2_〉 was only 0.0058% (Fig. S1B). Since the sequence differences between the two QCT pools is much higher than the QCT and gene sequence differences, we expect ∼0.006% to be an upper bound on the amplification difference between QCTs and the gene. Introducing distinct sets of QCT pools into all reactions provides an internally controlled upper bound for the error of 〈*D_QCT_*〉 measurements, which can be important for detecting assay degradation due to low sample quality (e.g. ethanol or salt carryover from DNA extraction).

We then sought to quantify the amount of human DNA present in a sample using QCTs. The number of DNA molecules corresponding to the gene of interest was calculated as 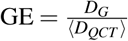; where GE is the haploid genome equivalents, and *D_G_* is the read depth of the gene (Fig. 2B). To more closely mimic the fragmentation pattern of cfDNA, human DNA was acoustically sheared to a mean fragment length of ∼150bp. A dilution series of 2-36ng sheared DNA was prepared with approximately 200 QCT molecules per reaction (Fig. S1C). PCR primers targeting a 150bp region of *HBB* exon 1 were used to co-amplify the QCT and sheared DNA molecules to yield a 150bp amplicon. Since not all sheared DNA molecules span both primer binding sequences, only a fraction of the sheared DNA would be amplifiable by PCR. After counting the number of amplifiable sheared DNA molecules, regression analysis showed that 64 GE/ng of sheared DNA was detectable, compared to the theoretical maximum of 278 GE/ng implied by a haploid genome mass of 3.6pg (Fig. S1C). This demonstrates the importance of directly measuring assayable genomic equivalents as opposed to DNA mass alone, since cfDNA is also fragmented with mean length ∼167bp.^29^ We then established the precision of GE estimates by measuring a smaller range of 30-33ng of sheared DNA with QCT molecules. Again, we reproducibly obtained genomic equivalents of ∼60GE/ng sheared DNA (Fig. 2E). Measured GE increased linearly with more DNA mass, increasing from 1850GE at 30ng to 2070GE at 33ng. The coefficient of variation (CV) of *HBB* molecular counts was 4.8%. Since the overall CV is affected by pipetting precision and Poisson sampling of limited DNA molecules (2.2% CV at 2000 molecules), 4.8% CV is the upper bound for QCT molecular counting precision.

### Ultra-rare variant calls enabled by QCTs

A particular challenge in detecting rare variants is false-positives due to contamination from another positive sample. This becomes particularly acute when multiple libraries are processed in parallel and sequenced in the same run.^30, 31^ Since the sequence diversity of QCT molecules aliquoted into each reaction is very small compared to the total QCT diversity in each synthesized pool, we realized that the set of EMI cluster sequences associated with each PCR can be used as a fingerprint to identify the sample. Given a total pool diversity of 1 million sequences, ∼10^443^ fingerprints are possible if 100 sequences are added to each PCR. Analysis of PCR fingerprints across the entire sequencing workflow can then be used to rule out the possibility that detected rare variants in any given reaction are from sample cross-contamination (Fig. 3).

**Figure 3.**
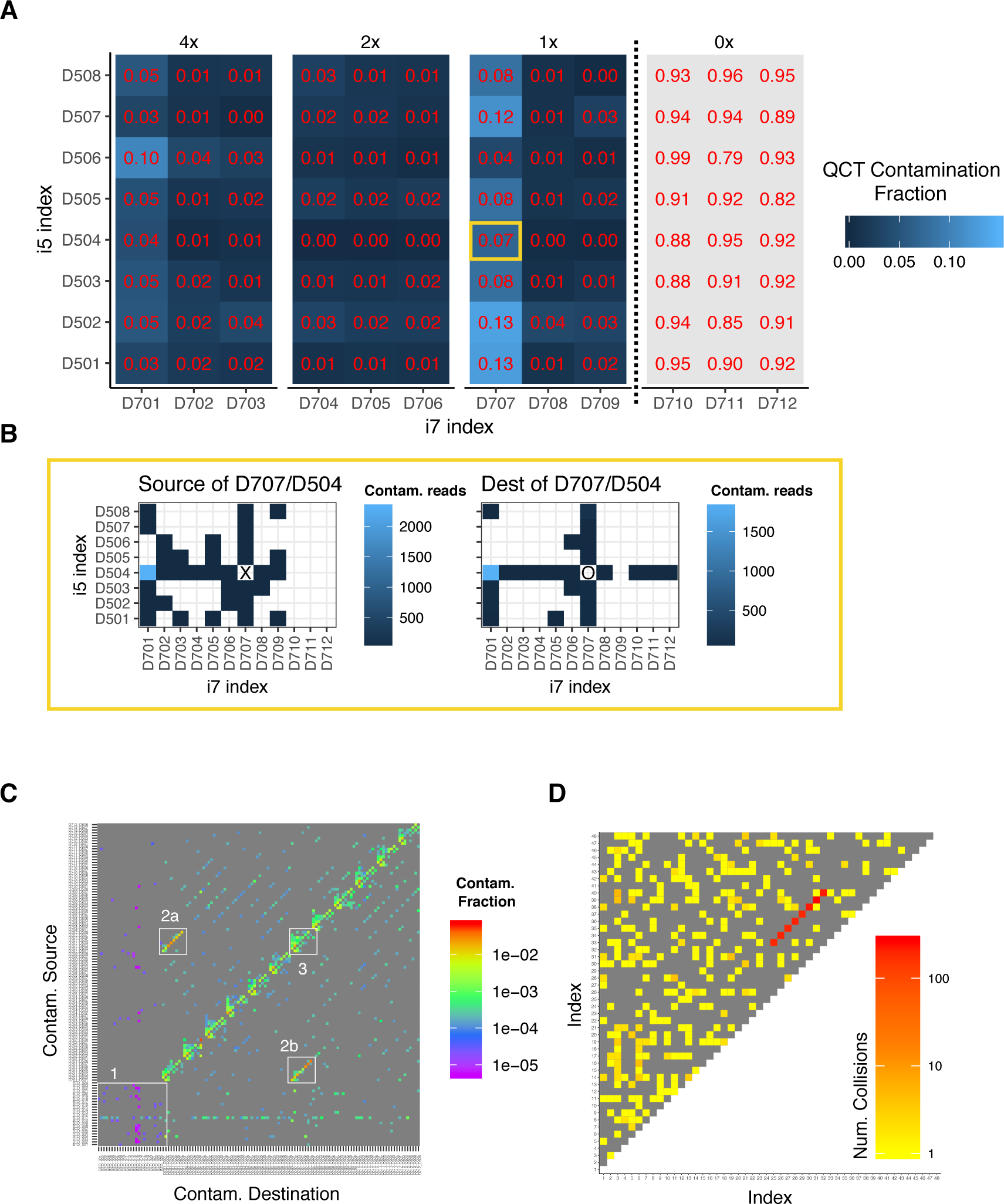
QCT tracking analysis quantifies contamination and sample mixups. Either 4x, 2x, 1x or 0 QCT molecules were added to 96 PCR reactions and sequenced on an Illumina Miseq lane. Within each PCR reaction, EMI sequence clusters were classified as contamination based on a read depth threshold (see Methods and Fig. S3). (A) The QCT contamination fraction for a given PCR is the proportion of QCT reads in that PCR well that are due to contamination from another PCR well. Wells are shaded and annotated as the fraction of its reads that are due to contamination. (B) Contamination can further be traced to specific PCR wells. Contaminating EMI clusters found in D707/D504 (yellow box, A) are traced to high-depth EMI clusters found elsewhere in the experiment (left), and the high-depth EMI clusters of D707/D504 are also found as contaminating reads in other wells (right). (C) Dual-unique indexes drastically reduce index misassignment. Both dual-unique indexes and Truseq-style barcoding were used on 120 samples prepared and sequenced in batch. Contamination source/destination is shown for all pairs of the 120 PCR reactions. The reactions that used unique dual indexes (1) had nearly 100-fold less contamination compared to Truseq-style barcoding. Cross-contamination of D701 and D707 is found at rates exceeding 1% (2a and 2b). Index misassignment between D5xx barcodes are also evident (3). (D) Pairwise analysis of QCT fingerprints identifies sample mixups. Forty-eight PCR reactions were processed in parallel, dual unique indexed, and sequenced on a Miseq lane. The similarity of QCT fingerprints is quantified as the number of high read depth EMI clusters in common (i.e. collisions) between two reactions. The number of collisions for all pairs of PCR reactions is shown.

We further analyzed the experiment performed in Fig. 2C to quantify contamination using QCT fingerprints. The sequencing depth for each EMI cluster in a PCR library was distinctly bimodal, with most EMI sequence clusters read at depth >30x. A minority of EMI sequencing reads were also present at depth 1-2x (Fig. S2). EMI sequencing clusters were classified as high-depth or low-depth (see methods). Low-depth EMI sequences could arise from (i) sequencing error, (ii) errors introduced during PCR amplification, (iii) cross-contamination during sample handling, or (iv) index misassignment.^30^ A low-depth EMI sequence cluster (typically 1-2x) observed in a PCR was classified as a contaminant if it was also observed at high-depth in a different PCR. We then computed the contamination fraction for each PCR as the number of contaminated QCT reads over total QCT reads (Fig. 3A). The PCRs with 0x QCT molecules should therefore register 100% contamination. QCT contamination analysis measured >90% contamination in 0x QCT wells, suggesting that this method is highly sensitive for detecting contamination. The remaining ∼10% of undetected contamination could be due to sequencing error or contamination that had occurred prior to PCR amplification. Unexpectedly, we found that PCR libraries barcoded with D701 and D707 indexes had high levels of total contamination consistently >5% and as high as 13% (Fig. 3A). A more granular analysis of contamination that traced contamination sources on a per-tube level revealed that for the reaction indexed by D707/D504, nearly all of the contamination originated from the D701/D504 reaction. Conversely, the D701/D504 well was the main destination of contaminating EMIs that originated from the D707/D504 reaction (Fig. 3B). Similar cross-contamination patterns were observed in the other wells indexed by D701 and D707. These data suggested to us that our D701 and D707 indexes themselves had become cross-contaminated, perhaps during oligo synthesis or index preparation. We also observed that most of the remaining contamination occurred in wells that have a D7xx or D5xx index in common, which is consistent with contamination due to index misassignment (Fig. 3C). To address these issues, we performed library barcoding with dual unique indexes.^32^ Tru-seq HT style combinatorial indexing using D7xx/D5xx pairs resulted in a median contamination of 0.5% (maximum 8.9%). Dual unique indexes reduced the observed contamination to 0.006% (maximum 0.03%), a nearly 100-fold improvement (Figs. 3C and S3).

In addition to contamination quantification, QCT analysis is also sensitive to sample mixups due to operator error during library preparation barcoding. If the same QCT fingerprint is observed in multiple samples at high read depth, this would indicate that a single assay was indexed to multiple barcodes. To quantify this, a sample collision score is defined as the number of high read-depth EMI clusters shared between two PCRs. We measured the number of colliding EMI sequences across all pairs of PCRs in an experiment where 8 reactions were barcoded twice in the same sequencing run. When the pairwise collisions are plotted, barcodes 25-32 immediately stand out because they share ∼150 EMI clusters in common with corresponding barcodes 33-40. This approach can therefore be used to identify common operator errors that would result in a reduced number of fingerprints in the sequencing data (Fig. 3D).

### Single-gene NIPT enabled by single base pair molecular counting

We next designed PCR assays for amplifying regions responsible for the most common genotypes of sickle cell disease, cystic fibrosis, spinal muscular atrophy (SMA), and alpha-thalassemia (Figs. 4 and S4). Sickle cell disease is most commonly caused by the rs334 (HbS) and rs33930165 (HbC) variants within exon 1 of *HBB*; over 90% of sickle cell disease cases are either HbSS or HbSC^33^. Since the paralogous genes *HBB* and *HBD* are highly similar in this region, >2bp of variation between *HBB* and *HBD* was included in the targeted region to ensure accurate read mapping (Fig. 4A). Cystic fibrosis is most commonly caused by recessive inheritance of rs113993960 (ΔF508, Fig. 4B) in *CFTR* exon 11. Statistical modeling of the accuracy of NIPT results uses the pathogenic allele fraction (AF) observed from maternal blood, the number of DNA molecules assayed for the allele fraction measurement, and the fetal fraction measurement (Fig. 4C).^34, 35^ Since SMA and alpha-thalassemia are caused by deletions, modified designs were used to quantify the pathogenic allele fraction (Fig. S4).

**Figure 4.**
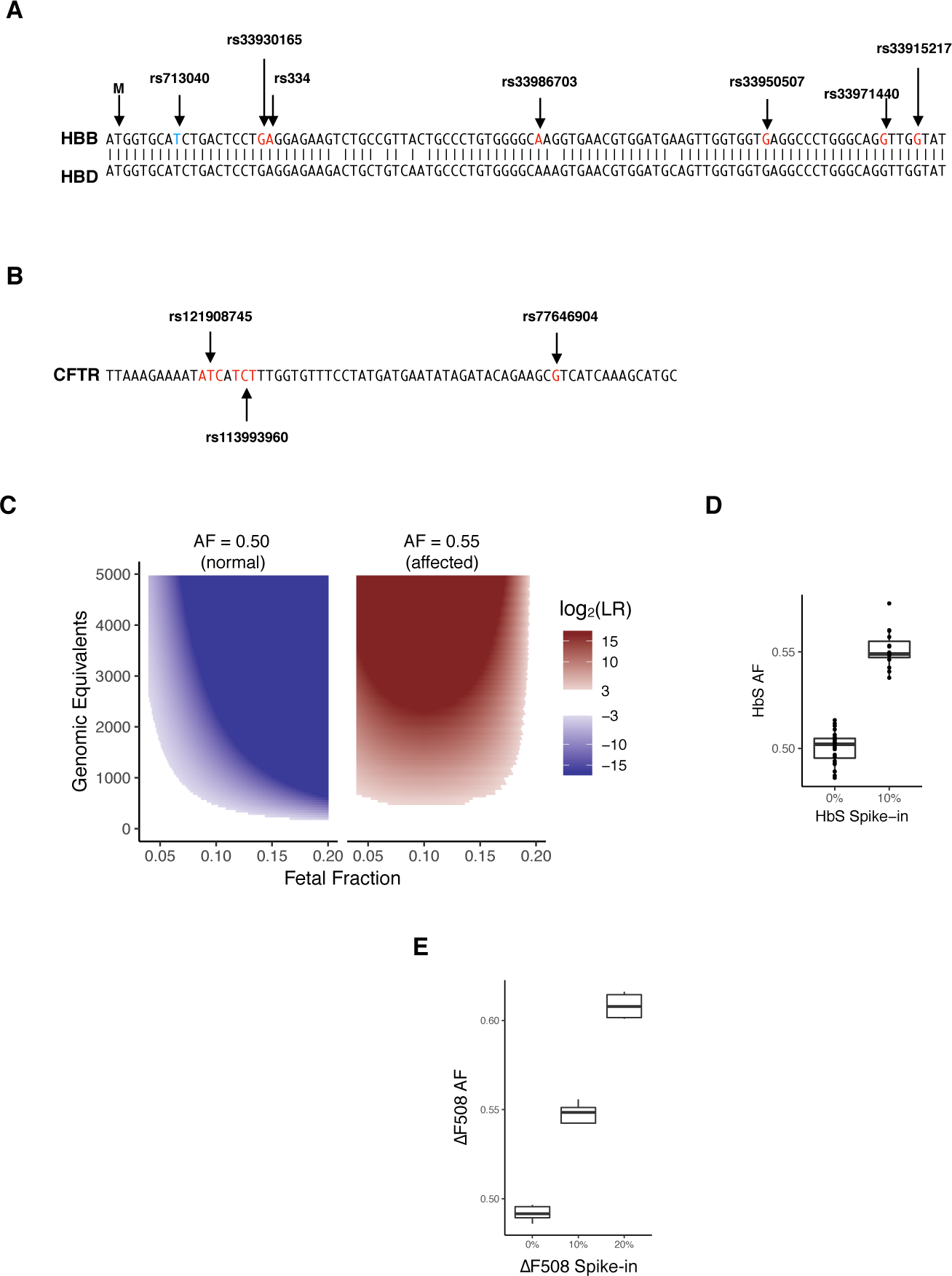
NIPT for sickle cell disease and cystic fibrosis by amplicon NGS. (A) Target region of *HBB* exon 1 NIPT and alignment with *HBD*. Start codon is shown as ‘M’. rs713040 is a benign variant with high allele frequency. rs33930165 and rs334 are pathogenic variants. (B) Target region for CFTR NIPT of the ΔF508 variant. (C) The NIPT assay uses pathogenic allele fraction, fetal fraction, and assayed genomic equivalents data to compute a likelihood ratio (LR) for an affected homozygous vs heterozygous fetus. Regions where 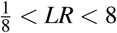 are shaded white to represent no-call regions. Affected NIPT calls are shaded blue, and non-affected calls are shaded red. (D) Validation of allele fraction measurement for sickle cell disease. Genomic DNA corresponding to a heterozygous sickle cell (NA16266) mother was sheared to 150bp and 0 to 10%sheared sickle cell disease DNA (NA16265) was mixed in. The HbS allele fraction (AF) was measured by amplicon NGS sequencing. When no SCD DNA is mixed in, the mean HbS AF = 0.500 (n=28, CV=1.6%). With 10% SCD DNA, mean HbS AF = 0.551 (n=15, cv=1.7%). QCT analysis measured 2000GE in these assays. (E) Validation of allele fraction measurement for cystic fibrosis. Sheared homozygous cystic fibrosis variant ΔF508 was spiked-in to 30ng heterozygous ΔF508 sheared DNA at the given proportions.

We first validated that accurate allele fraction measurements could be obtained from *HBB* exon 1. We generated ersatz cfDNA samples by shearing genomic sickle cell DNA to ∼150bp. Approximately 33ng total of sheared DNA was added to each PCR assay, and QCT analysis showed that 2,000 GE of *HBB* exon 1 were amplified from sheared DNA. In the ersatz samples of heterozygous sickle cell DNA (NA16266), we measured a mean HbS allele fraction of 0.500 +/- 0.0015 standard error of the mean (SEM) across 28 PCR replicates (Fig. 4D), indicating that *HBB* alleles are amplified without introducing any bias. The CV of the replicates was 1.3%-2.2% (95% confidence interval). Importantly, the CV is in accordance with the Poisson noise of 2.2% associated with 2000 molecules, suggesting that the assay for *HBB* allele fraction is operating at the physical limit of counting statistics.

We next prepared a sheared DNA mixture of 90% heterozygous DNA and 10% homozygous sickle cell disease DNA to mimic cfDNA from a sickle cell carrier pregnancy with 10% SCD fetal fraction. The measured HbS allele fraction matched very closely with the expected HbS allele fraction of 0.55 (0.549-0.554, 95% CI; n=15). The large separation between the 0% and 10% SCD mixtures shows that pathogenic allele fraction differences for a sickle cell trait (heterozygous) vs sickle cell disease (homozygous) fetus can be easily discriminated. Similar results were obtained for >200 PCR replicates at mixtures ranging from 0% to 20% for HBB, cystic fibrosis, SMA, and alpha-thalassemia (Figs. 4E and S4).

To confirm that sheared DNA behaves similarly to cell-free DNA, we next obtained 30 blood samples from male and female patients that are compound heterozygotes for sickle cell disease at Baylor College of Medicine (Fig. 5). cfDNA was purified from blood plasma and the *HBB* allele fraction assay was performed. The results of the *HBB* allele fraction assay again agreed with the expected ^1^ allele fraction (Fig. 5A). On average, 3500GE of *HBB* were assayed for each 10mL blood sample. We measured 2.2% CV (1.6-2.9% 95% CI), which was in good agreement with the expected CV=1.7% associated with Poisson counting of 3500GE. The proportion of *HBB* DNA amenable to PCR amplification was 129GE/ng cfDNA (Fig. 5B), which is almost twice what we previously observed for sheared DNA (Fig. 2D). The increased numbers of assayable GE from cfDNA shows that the analytical validation performed with ersatz samples was more challenging than for cfDNA blood samples. This difference in capture efficiency could be because acoustically sheared DNA (150bp) was more fragmented than cfDNA (165bp) and/or differences between acoustic shearing and biological mechanisms of cfDNA fragmentation.^29^ Since none of the blood samples were taken from pregnancies, they can also serve as negative controls for NIPT analysis. None of the allele fraction measurements in these non-pregnant controls would have resulted in positive NIPT results (Fig. 5A).

**Figure 5.**
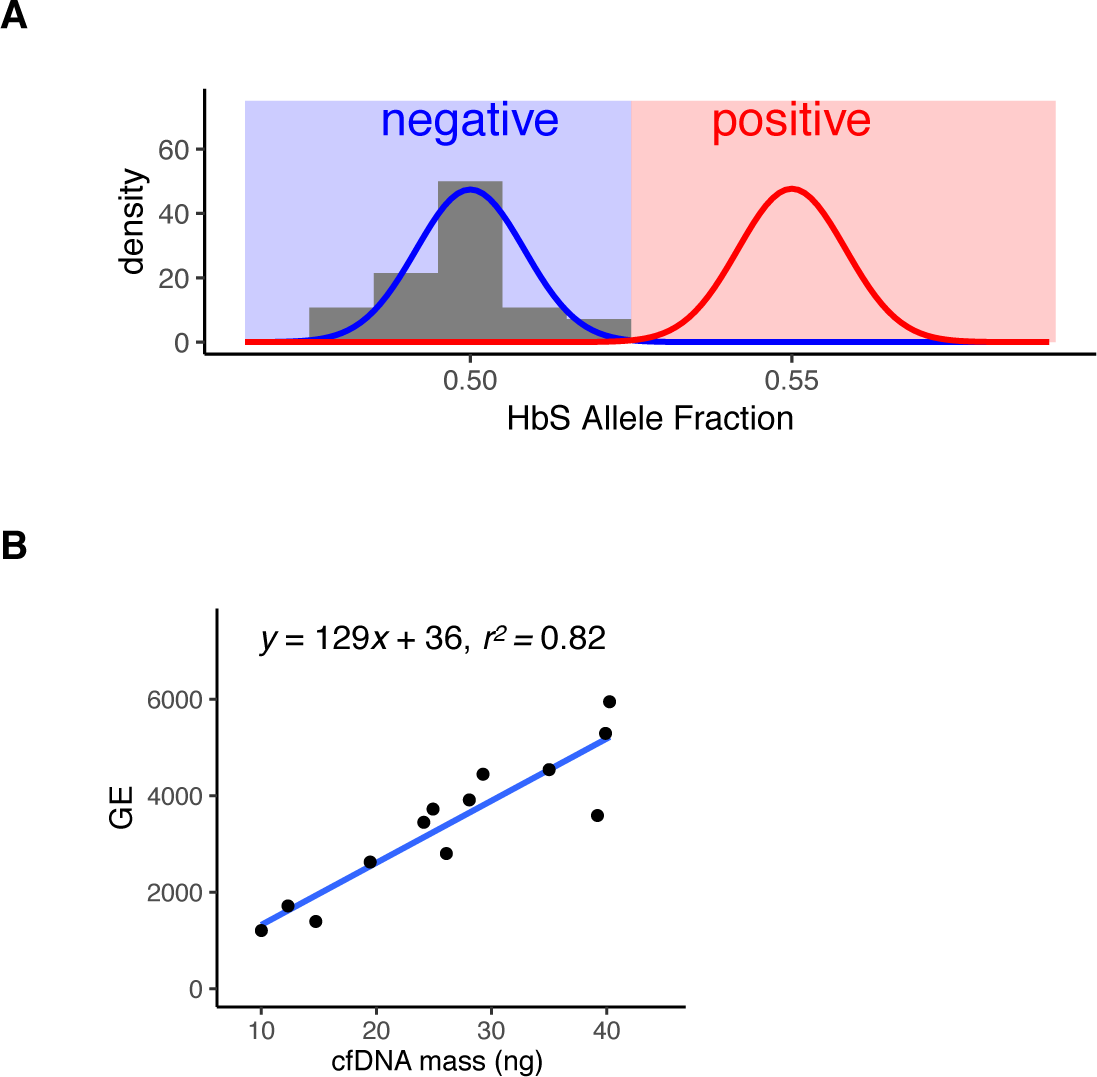
SCD NIPT on negative control cfDNA resulted in no false-positives. Optimized *HBB* probes were validated for use on cfDNA by performing the NIPT assay on 10mL of venous blood from non-pregnant compound heterozygotes (HbAS or HbSC). The mean HbS AF was 0.498 with a coefficient of variation of 2.2% (n=30). The mean number of *HBB* molecules assayed in each blood tube was 3500. Our results agree with the expected HbS AF=1/2 and CV=1.7%; the histogram of measured HbS AF corresponds very well with the theoretical binomial distribution in blue (AF=1/2, n=3500). Assuming that positive cases have a 10% fetal fraction (red curve), none of the negative controls would have been called as positive for fetal SCD. (B) Assayed genomic equivalents of *HBB* exon 1 in cfDNA. The concentration of 13 cfDNA samples was quantified by Qubit to determine the mass of cfDNA used in the *HBB* assay. On average, 1 ng of cfDNA resulted in the capture of 129 haploid genomic equivalents of *HBB*.

### Clinical validation of single-gene NIPT

We next obtained maternal blood and saliva samples from 208 healthy pregnant donors from Yashoda Hospital, Ghaziabad, India. Ethical clearance was obtained from the Yashoda Institutional Ethics Committee (IEC: ECR/970/Inst/UP/2017). While beta-thalassemia is common in India with a carrier rate as high as 5%, the distribution of pathogenic alleles is heterogeneous across subpopulations and geographic areas^36^. However, when we genotyped these samples, there were only 3 pregnant donors who were beta-thalassemia carriers. Therefore, to clinically validate *HBB* NIPT by molecular counting, we performed NIPT analysis for a linked, benign variant that is located within the target region of *HBB* exon 1 (Fig. 4A). Since *HBB* NIPT is an amplicon-based NGS assay, the ability to detect fetal inheritance of this variant (rs713040) is identical to that of SCD or any other pathogenic variants that are found in the target region of exon 1 (e.g., HbS, HbC, HbE, IVS1,1, IVS1,5). Moreover, because this variant has an extremely high minor allele frequency (0.2), we were able to obtain all maternal-fetal genotype combinations that represent healthy and disease states.

Follow-up *HBB* genotyping of newborns was obtained for 52 pregnancies. However, the amount of cfDNA recovered from corresponding maternal plasma samples using the *HBB* allele fraction assay was found to be abnormally low (mean GE=996, min GE=21) compared to the cfDNA purified from negative controls collected at Baylor College of Medicine (mean GE=3500, min GE=1200; Fig. 5). Fifteen of the pregnant cfDNA samples contained <200 GE of assayed *HBB*. Since 200 GE corresponds to <2ng of cfDNA, these 15 samples were excluded from further analysis. Previous reports have shown that ∼4000GE should be recoverable from a 10mL blood sample.^34, 37^ The paucity of cfDNA recovered from these samples may be explained by incomplete filling of blood tubes, sample loss during cell-free DNA purification, or DNA degradation during extended storage at −20C (5-8 months).

Our approach for NIPT integrates measurements of (i) fraction of fetal DNA present in cfDNA, (ii) molecular counts of assayed cfDNA, (iii) allele fraction of the maternal variant, and (iv) allele fraction of any variants that are not present in the maternal genotype (distinct paternally inherited variants). Approximately 1/3 of each purified cfDNA sample was used to determine the fetal fraction using a custom amplicon NGS assay that interrogates 86 common SNVs across all autosomal chromosomes (Fig. S5). QCT molecules were added to the remaining cfDNA and the *HBB* allele fraction assay was performed to determine molecular counts and allele fractions of *HBB* exon 1 variants. These measurements were then used in a statistical model to determine the genotype of the fetus. Because a fetus inherits one allele from each parent, in cases where the maternal genotype is homozygous, the detection of any non-maternal allele indicates that the fetus is heterozygous for that variant, i.e., inherited a paternal allele (paternal inheritance).

On the other hand, for a sample where the maternal genotype is heterozygous, NIPT for recessive inheritance requires precise quantification of allele fraction to determine whether fetal alleles are contributing to an observed allele fraction significantly different from 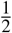. For example, the expected variant allele fraction (VAF) in cfDNA for a heterozygous fetus will remain at 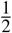, the same as the maternal level. However, if the fetus is homozygous for the variant, i.e., the fetus has inherited two identical alleles with the pathogenic variant, the VAF from the cfDNA sample should increase to 0.55 for a sample with 10% fetal fraction, to 0.60 VAF for 20% fetal fraction, and so on (recessive inheritance). The probability distributions of these allele fractions depends on the molecular counts due to Poisson counting noise (Figure 4C).

In samples that had a homozygous (C/C reference or T/T variant) maternal genotype for the benign variant rs713040, we performed *HBB* NIPT by detecting a distinct paternally inherited fetal allele (n=14; Table 1). In these pregnancies, a heterozygous fetal genotype can occur only when the fetus inherits a paternal allele different from the maternal allele, and the resultant minor allele fraction (MAF) in cfDNA is expected to be 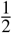 of the fetal fraction (e.g. MAF of 0.05 when fetal fraction is 10%). Likewise, a homozygous fetus matching the genotype of the mother is determined by the absence of a paternal allele. This detection requires distinguishing a true, rare variant, i.e., the paternal allele, from any other sources of low-frequency variants introduced either by contamination, index misassignment, amplification, and/or sequencing errors. For these samples, QCT analysis indicated sample contamination and index misassignment was negligible (median=0.01%, max=0.2%). Then, we used a statistical model to ensure any observed minor alleles could not be explained by sequencing/polymerase error (Fig. 6A-B; discussed below). Since QCT analysis showed that these samples were sequenced to >50 reads/molecule, the contribution of read depth to noise is negligible, and sampling noise is nearly entirely due to the number of DNA molecules assayed. To rule out the possibility that the observed minor allele molecules are present due to sequencing error, we calculated two probability distributions for each sample: one for background sequencing error and the second for a paternally inherited allele. The fetal genotype is then determined by which probability distribution more closely matches the observed result. Hypothetically, the measured allele fraction could occur where the probability distributions have overlap. The likelihood ratio (LR) roughly corresponds to how likely it is for the observed measurement to be positive or negative, with LR > 1 indicating increased evidence for a positive result (affected fetus). To avoid calling NIPT results on inconclusive evidence, a no-call threshold was set at 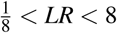, a standard that has previously been used in single-gene NIPT^34^. Examples of this analysis are shown in Figs. 6A and 6B for the cases of a homozygous and heterozygous fetus, respectively. The LR for all 14 paternal inheritance cases exceeded 10,000-fold, and *HBB* genotyping of newborns resulting from these pregnancies confirmed that all of the NIPT calls were correct. This was true even in highly challenging samples with limited fetal fraction (<5%) or quantity of cfDNA available (<250GE). Out of the 14 maternal blood samples analyzed for paternal inheritance, 2 were homozygous negative C/C, 2 were heterozygous negative C/T, and 10 were homozygous positive T/T (Table 1).

**Figure 6.**
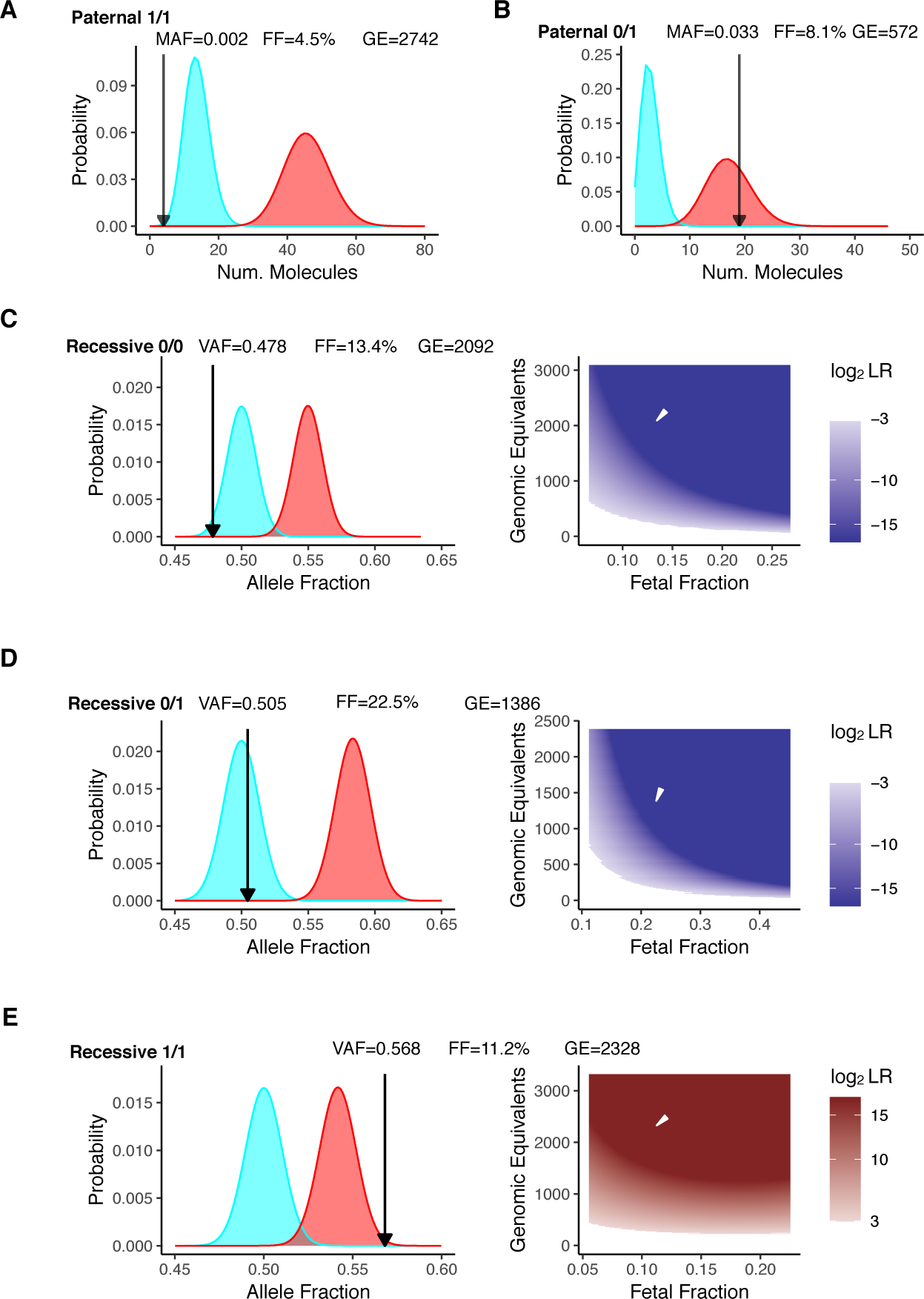
NIPT of rs713040 in *HBB* exon 1. (A) Probability distribution of sample 04B for paternal inheritance resulting in a NIPT homozygous call. (B) Probability distribution of sample 38C for paternal inheritance resulting in a NIPT heterozygous call. Black and white arrows indicate diagnostic measurements of allele fraction, GE, and fetal fraction. (C) Negative call for recessive NIPT resulting in a homozygous child. (D) Negative recessive NIPT call resulting in a heterozygous child. (E) Positive recessive NIPT call resulting in an ‘affected’ child.

**Table 1.**
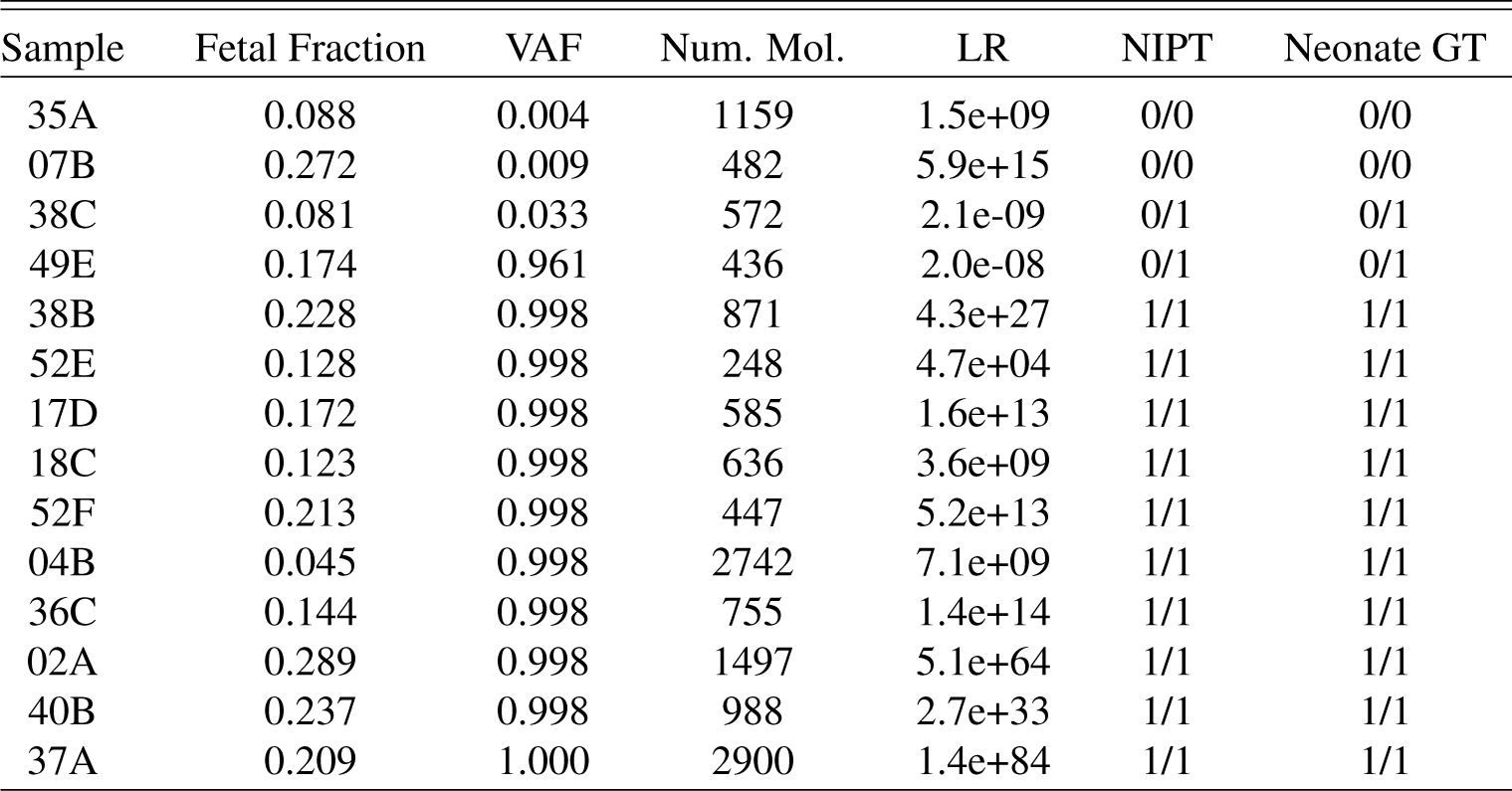
NIPT for paternal inheritance of rs713040

| Sample | Fetal Fraction | VAF | Num. Mol. | LR | NIPT | Neonate GT |
| --- | --- | --- | --- | --- | --- | --- |
| 35A | 0.088 | 0.004 | 1159 | 1.5e+09 | 0/0 | 0/0 |
| 07B | 0.272 | 0.009 | 482 | 5.9e+15 | 0/0 | 0/0 |
| 38C | 0.081 | 0.033 | 572 | 2.1e-09 | 0/1 | 0/1 |
| 49E | 0.174 | 0.961 | 436 | 2.0e-08 | 0/1 | 0/1 |
| 38B | 0.228 | 0.998 | 871 | 4.3e+27 | 1/1 | 1/1 |
| 52E | 0.128 | 0.998 | 248 | 4.7e+04 | 1/1 | 1/1 |
| 17D | 0.172 | 0.998 | 585 | 1.6e+13 | 1/1 | 1/1 |
| 18C | 0.123 | 0.998 | 636 | 3.6e+09 | 1/1 | 1/1 |
| 52F | 0.213 | 0.998 | 447 | 5.2e+13 | 1/1 | 1/1 |
| 04B | 0.045 | 0.998 | 2742 | 7.1e+09 | 1/1 | 1/1 |
| 36C | 0.144 | 0.998 | 755 | 1.4e+14 | 1/1 | 1/1 |
| 02A | 0.289 | 0.998 | 1497 | 5.1e+64 | 1/1 | 1/1 |
| 40B | 0.237 | 0.998 | 988 | 2.7e+33 | 1/1 | 1/1 |
| 37A | 0.209 | 1.000 | 2900 | 1.4e+84 | 1/1 | 1/1 |

NIPT for recessive inheritance of *HBB* was then performed on samples with the heterozygous rs713040 maternal genotype, T/C. In these samples, the fetal genotype could be (i) homozygous reference allele, T/T; (ii) heterozygous T/C; or, (iii) homozygous for variant allele C/C. To more closely match the clinical use case of determining fetal disease status for a recessive disorder, the T/T and T/C genotypes were considered ‘normal,’ while the C/C was considered ‘affected.’ As in paternal inheritance NIPT, recessive inheritance NIPT uses a statistical model to compute the likelihood ratio of an affected vs. normal fetal genotype. Out of the 13 samples in which NIPT results and newborn follow-up were available, 5 were C/C affected, 3 were T/T normal, and 5 were T/C normal (Table 2, Fig. 6C-E). All NIPT results agreed with follow-up neonatal genotyping. Recessive inheritance NIPT analysis was performed on an additional 10 cfDNA samples, but the results were indeterminate based on the LR thresholds (Table S1). Notably, even lower amounts of cfDNA were recovered from the no-call samples (mean GE=600) compared to the samples for which recessively inherited NIPT calls were made (mean GE=1800). The strength of incorporating molecular counting into NIPT is that unreliable samples can be identified and incorrect calls can be avoided. Given that cfDNA obtained from *HBB* heterozygotes at Baylor College of Medicine had mean cfDNA molecular counts of 3500GE (Fig. 5), we expect the no-call rate to not be an important issue in a clinical setting. Overall, 27/27 NIPT calls, including paternal and recessive inheritance, were concordant with the newborn *HBB* genotype.

**Table 2.** NIPT for recessive inheritance of rs713040

| Sample | Fetal Fraction | VAF | Num. Mol. | LR | NIPT | Neonate GT |
| --- | --- | --- | --- | --- | --- | --- |
| 30B | 0.153 | 0.477 | 8616 | 1.0e-44 | normal | 0/0 |
| 31B | 0.171 | 0.478 | 304 | 1.5e-02 | normal | 0/0 |
| 08A | 0.134 | 0.478 | 2092 | 3.5e-09 | normal | 0/0 |
| 56C | 0.325 | 0.482 | 663 | 7.2e-12 | normal | 0/1 |
| 39A | 0.079 | 0.498 | 1868 | 2.3e-02 | normal | 0/1 |
| 63B | 0.225 | 0.505 | 1386 | 2.4e-08 | normal | 0/1 |
| 47F | 0.308 | 0.508 | 1319 | 6.2e-14 | normal | 0/1 |
| 09B | 0.208 | 0.521 | 1601 | 1.3e-04 | normal | 0/1 |
| 38A | 0.112 | 0.568 | 2328 | 7.9e+07 | affected | 1/1 |
| 25B | 0.242 | 0.569 | 360 | 1.7e+01 | affected | 1/1 |
| 40E | 0.184 | 0.570 | 573 | 2.1e+02 | affected | 1/1 |
| 50C | 0.179 | 0.586 | 1085 | 3.4e+06 | affected | 1/1 |
| 17B | 0.219 | 0.603 | 1611 | 1.8e+14 | affected | 1/1 |

## Discussion

There is significant unmet medical need for NIPT of these recessively inherited single-gene disorders. Affected individuals face significantly reduced life-span and require frequent access to intensive medical care. Current medical guidelines set by the American College of Obstetrician & Gynecologists and American College of Medical Genetics recommend that all pregnancies should be screened to determine the carrier status of the parents before offering genetic counseling and invasive diagnostic procedures.^21^ Moreover, reducing the need for paternal DNA could be a significant improvement in clinical practice. Current screens that test the mother for carrier status result in 11% to 30% positive calls and require follow-up with paternal carrier testing.^38^ However, paternal carrier testing may be unavailable or unreliable. Paternity is not established at the time of birth in 30% of US births to single-mothers,^39^ and this is in addition to >0.8% hidden non-paternity that has been uncovered in population-level genetic studies.^40^

Single-gene NIPTs require precision at the level of molecular counts. Digital PCR is an alternative method for obtaining molecular count data.^19, 20^ In fact, NIPT for sickle cell disease and beta-thalassemia has previously been demonstrated in proof-of-principle studies using dPCR.^41, 42^ However, dPCR has two critical limitations that hinder its translation to the clinic. First, dPCR relies on ∼15-30nt SNV-specific probes and therefore does not the have the single-base pair resolution of a sequencing approach. If nearby cis-variants are present within this footprint or if a high homology region is also amplified, a significant proportion of cases can be missed by dPCR. For instance, this is particularly acute in sickle cell disease, where HbC, HbS, and rs713040 are all within 12bp of each other; and where the *HBD* gene is a close paralog of *HBB*. These factors might have contributed to the 80% accuracy obtained in a previous study that used dPCR for NIPT of sickle cell disease.^41^ In addition, the multiplexability of digital PCR is limited to only 2-4 variants. This significantly limits its throughput and use for multiple variants, disorders, or samples at the same time, and has become a crucial impediment to its widespread clinical use.

To overcome these limitations, we developed a method for counting DNA molecules using amplicon-based next-generation DNA sequencing and QCT analysis. The QCT molecular counting approach uses synthetic DNA that co-amplifies with the gene of interest (e.g. *HBB*, *CFTR*, etc.) to serve as a reference standard. The exact number of QCT molecules added to each sample is reconstructed from sequencing data which then enables molecular count information of the gene of interest to be recovered from sequencing depth data. Therefore, the synthetic QCT DNA used as a reference standard is calibration free and not susceptible to the Poisson noise associated with addition of ∼100-400 synthetic DNA molecules. Furthermore, since QCT molecular counting is compatible with amplicon sequencing, multiple loci can be simultaneously interrogated via multiplex PCR, and as many as 50-100 sgNIPT assays can be pooled and sequenced on a single Miseq lane (200,000-400,000 reads per assay). Single-gene NIPTs using QCTs were developed for sickle cell disease, cystic fibrosis, beta-thalassemia, alpha-thalassemia, and spinal muscular atrophy. *HBB* NIPT was performed on blood taken from pregnant women, and results were confirmed by follow-up DNA sequencing of newborns, resulting in 100% concordant NIPT calls. Molecular counting ability enables statistical modeling of NIPT results so that false calls can be avoided with even the most challenging samples.

In addition to improvements in single-base pair resolution and multiplexability, QCT molecular counting has 25-100x more dynamic range than digital PCR. Droplet digital PCR is most precise at measuring ∼20,000 molecules, with an upper limit of 100,000 molecules.^19^ Even low-throughput NGS instruments such as the Illumina MiSeq produce 25 million reads, which can be used to count 2.5 million molecules using the QCT method without any decrease in accuracy. This increase in dynamic range unlocks absolute quantification of DNA and RNA levels in a diverse range of applications including T-cell receptor profiling, transcriptomics, and microbiome applications.

Prior to QCT molecular counting, diverse adapter/primer pools called UMIs have been used for detection of unique molecules as well as low resolution molecular counting.^17, 43^ QCTs rely on a similar computational approach in which sequence diversity is used to identify single molecules. However, the QCT approach embeds sequence diversity within the target region instead of requiring a ligation or pre-amplification step inherent to UMIs. This has several advantages, including (i) it avoids inefficient ligation and pre-amplification PCR steps that can result in >90% loss of the sample, resulting in lower sensitivity; and sequence diversity is contained within ∼100 molecules in the QCT approach, as opposed to the picomole (10^11^ molecules) amounts of UMIs that can lead to primer/adapter dimers and hinder multiplexing.^44, 45^

We also found that QCTs were useful for ensuring the integrity of sample preparation and sequencing in addition to molecular counting. We point out that these features should also enable more reliable and sensitive rare variant detection in liquid biopsy. Currently, the limit of detection (LOD) for circulating tumor DNA is typically given in terms of allele fraction. However, an LOD of 0.1% allele fraction is not meaningful when a particular sample only contains 500 haploid genomic equivalents (less than 1 molecule). The QCT molecular counting approach enables an LOD to be calculated as an absolute number of tumor DNA molecules. Furthermore, reliable rare-variant detection requires assurances that minor alleles are not the result of cross-contamination from previously processed or positive control samples. We have integrated QCT cross-contamination analysis into our workflow for detecting contamination from fluid handling, operator error, and index misassignment. Measurements of contamination and index misassignment in every sample, rather than only as a negative control, enable a lower limit of detection to be obtained in liquid biopsy applications. Since QCTs are internal amplification controls that are present in every sample, they are also sensitive to and can help identify common problems that can negatively impact assay performance, such as PCR inhibition from hemolysis or salt and ethanol carryover during DNA purification.

Perhaps, the highest technical impact of QCT molecular counting may be on quantification of gene copy number variation (CNV). 5-10% of the human genome consists of CNV >50bp, and an additional 33% of the genome is susceptible to CNV in cancerous tissues.^46, 47^ While CNVs are often seen as hallmarks of cancer, current liquid biopsy approaches are limited in their capability for CNV detection. Our NIPT results, particularly those that require copy number variation detection such as spinal muscular atrophy and alpha-thalassemia, suggest that QCT molecular counting can detect a single additional copy of a gene at 5% allele fraction. This is a significant improvement to current liquid biopsies that can only detect 6 additional copies or more of an oncogene, e.g., *HER2*, at tumor fractions of 5-20%.^14, 15^

## Methods

### QCT synthesis and analysis

QCT pools were synthesized by IDT as 4nmol scale ultramers. Sequences for *HBB* Exon 1 QCTs are given in Table S2. QCT oligos were double stranded by addition of primer hbs_qct_pext and incubation with Klenow polymerase at 37C for 1 hour. Double stranded QCT pools were then gel-purified and diluted to 15fM in TE-Tween (10mM Tris, 0.1mM EDTA, 0.05% Tween-20).

QCT reads matching the QCT identifier were extracted from raw sequencing reads. For each PCR reaction, EMI sequence clusters were generated by grouping EMI sequences together if they differ by 2 or fewer mismatches. EMI sequence clusters were then thresholded into high- or low- read depth clusters by read depth. The read depth threshold is computed for each PCR reaction as the square root of the mean EMI sequence cluster read depth. The number of QCT molecules in that PCR reaction is the number of high-depth EMI sequence clusters, and 〈*D_QCT_*〉 is the mean read depth per QCT molecule. Contaminating QCT reads are identified by a low-read depth EMI sequence cluster in one PCR assay that appears in another PCR assay at high-read depth. The EMI fingerprint for the PCR reaction is the set of high read depth EMI sequences. The assayed genomic equivalents of *HBB* exon 1 DNA is computed as 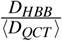, where *D_HBB_* is the read depth of the *HBB* Exon1 amplicon.

### DNA Purification and Library Prep

Approximately 10mL of venous blood was collected into a cfDNA blood collection tube (Streck, Omaha, NE). Plasma was separated from whole blood according to manufacturer instructions and stored at −20C until further processing. Cell-free DNA was purified from plasma via the Qiagen Circulating Nucleic Acid kit using an elution volume of 50ul.

An 86-plex amplicon NGS assay was designed to measure paternal inheritance of common allele frequency SNVs (Fig. S5). The fetal fraction was determined as 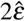, where 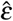 is the median MAF of paternally inherited alleles across at least 9 loci. 10ul of cfDNA was used per fetal fraction assay.

To perform the *HBB* exon 1 allele fraction assay, approximately 200 HBB QCT molecules were added to 35ul cfDNA, and the mixture was PCR-amplified and sequenced on the Illumina Miseq using dual unique indexes.

### Non-pregnant cell-free DNA controls

Samples were obtained from 30 pediatric patients with SCD receiving care at Texas Children’s Hospital Hematology Center. Samples were collected from both genders, aged 2-18 years of age, obtained under a Baylor College of Medicine Internal Review Board approved protocol. Venous blood was collected in 10mL Streck cfDNA tubes, and he cfDNA was purified from plasma using the Qiagen circulating nucleic acids kit. The HBB allele fraction assay was performed on all samples. The remaining cfDNA in these samples were either used in the fetal fraction assay as negative controls or analyzed by fluorometry (Qubit) to compare DNA mass and assayed genomic equivalents (Fig. 5B).

### Pregnant maternal blood samples

Buccal swabs and 6-10mL venous blood in Streck tubes were obtained from 208 pregnancies at >12 weeks of gestation at Yashoda Hospital, Ghaziabad, India. Ethical clearance for this study was obtained from the Institutional Ethics Committee of Yashoda Hosipital (IEC: ECR/970/Inst/UP/2017). Of these patients, we were able to obtain buccal swabs of 52 newborns for confirmation of NIPT results. *HBB* genotyping of the newborns was performed by PCR amplification of the *HBB* gene using primers HBBNextera500F1 and HBBNextera500R1 to generate a 2.5kb amplicon, followed by Nextera-based library preparation and Miseq sequencing. The GATK Hapolotype Caller was then used to call variants from sequencing data.^48^ Cell-free DNA was purified from maternal plasma by the Qiagen Circulating Nucleic Acids kit with an elution volume of 50ul. 10ul of cfDNA was used for measuring fetal fraction. 35ul of cfDNA was used for the *HBB* exon 1 allele fraction assay.

### NIPT Statistical Modeling

The binomial distribution was used to model the probability of measuring the *HBB* allele fraction, *x*; given fetal fraction, 2*ε*; and DNA molecule count, *N*. The likelihood of a paternal allele is then 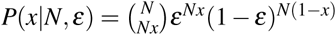, and the probability of sequencing error is 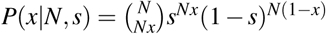. The rate of sequencing error, *s*, was set at 0.5%. The LR that the fetus inherited a paternal allele compared to sequencing error then the ratio of these two probabilities, 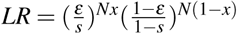. For maternal inheritance, we computed likelihoods for an affected vs heterozygous fetus, 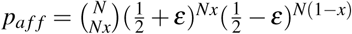 and 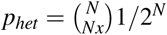, and samples that had likelihood ratio 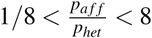 where considered no-calls. Fetal fraction of *HBB* exon 1 was adjusted by a factor of 0.74 to match levels observed in pregnant samples without follow-up (Fig. S4).

## Acknowledgements

We gratefully acknowledge Vimal Jain, Mukesh Bhardwaj, Dr. Amit Bhalla, Dr. Sanjay Srivastava, Dr. Pradeep Varshney, Dr. Shilpa Chapadgaonka for their assistance with sample collection at Yashoda Hospital and for providing laboratory facilities at Manav Rachna International University.

Research reported in this publication was supported by the National Heart, Lung, and Blood Institute of the National Institutes of Health under award number R43HL144322. 10% was funded by federal sources. 90% was funded by BillionToOne. The content is solely the responsibility of the authors and does not necessarily represent the official views of the National Institutes of Health.

## Author contributions statement

DST, SS, BPL, VAS, and OA conceived and designed experiments. DST, SS, BPL, AN, RS, NI, CKK, and OA conducted experiments. DST, BPL, and OA analyzed the results. DST and OA wrote the manuscript. All authors reviewed the manuscript.

## Additional information

DST, SS, BPL, AN, and OA are or were employees of BillionToOne and hold stock or options to hold stock in the company. Patent disclosures about this work have been filed with BillionToOne, Inc. BillionToOne, Inc. funded portions of this study.

## References

1. Broad. Broad Institute sequences its 100,000th whole human genome on National DNA Day.

2. Buenrostro, J. D., Wu, B., Chang, H. Y. & Greenleaf, W. J. ATAC-seq: A method for assaying chromatin accessibility genome-wide. Curr. Protoc. Mol. Biol. 2015, 21.29.1–21.29.9, DOI: 10.1002/0471142727.mb2129s109 (2015). 15334406.

3. Strobel, E. J., Yu, A. M. & Lucks, J. B. High-throughput determination of RNA structures, DOI: 10.1038/s41576-018-0034-x (2018).

4. Meng, L. et al. Use of exome sequencing for infants in intensive care units ascertainment of severe single-gene disorders and effect on medical management. JAMA Pediatr. 171, e173438, DOI: 10.1001/jamapediatrics.2017.3438 (2017).

5. Blumenthal, G. M. et al. Oncology Drug Approvals: Evaluating Endpoints and Evidence in an Era of Breakthrough Therapies. The Oncol. 22, 762–767, DOI: 10.1634/theoncologist.2017-0152 (2017).

6. Blumenthal, G. M. & Pazdur, R. Approvals in 2017: gene therapies and site-agnostic indications. Nat. Rev. Clin. Oncol. 15, 127–128, DOI: 10.1038/nrclinonc.2018.11 (2018).

7. Diaz, L. A. & Bardelli, A. Liquid biopsies: Genotyping circulating tumor DNA, DOI: 10.1200/JCO.2012.45.2011 (2014). 15334406.

8. Siravegna, G., Marsoni, S., Siena, S. & Bardelli, A. Integrating liquid biopsies into the management of cancer. Nat. reviews. Clin. oncology 14, 531–548, DOI: 10.1038/nrclinonc.2017.14 (2017). 15334406.

9. Fan, H. C., Blumenfeld, Y. J., Chitkara, U., Hudgins, L. & Quake, S. R. Noninvasive diagnosis of fetal aneuploidy by shotgun sequencing DNA from maternal blood. Proc. Natl. Acad. Sci. 105, 16266–16271, DOI: 10.1073/pnas.0808319105 (2008). 0705.1030.

10. Norton, M. E. et al. Cell-free DNA Analysis for Noninvasive Examination of Trisomy. New Engl. J. Medicine 372, 1589–1597, DOI: 10.1056/NEJMoa1407349 (2015).

11. van Schendel, R. V., van El, C. G., Pajkrt, E., Henneman, L. & Cornel, M. C. Implementing non-invasive prenatal testing for aneuploidy in a national healthcare system: global challenges and national solutions. BMC Heal. Serv. Res. 17, 670, DOI: 10.1186/s12913-017-2618-0 (2017).

12. Zill, O. A. et al. The landscape of actionable genomic alterations in cell-free circulating tumor DNA from 21,807 advanced cancer patients. Clin. Cancer Res. 24, 3528–3538, DOI: 10.1158/1078-0432.CCR-17-3837 (2018).

13. Shlien, A. & Malkin, D. Copy number variations and cancer, DOI: 10.1186/gm62 (2009). 1512.00567.

14. Allen, J. M. et al. Genomic Profiling of Circulating Tumor DNA in Relapsed EGFR-mutated Lung Adenocarcinoma Reveals an Acquired FGFR3-TACC3 Fusion. Clin. Lung Cancer 18, e219–e222, DOI: 10.1016/j.cllc.2016.12.006 (2017).

15. Vowles, J. et al. Abstract 5705: Analytical validation of Guardant360 v2.10. Cancer Res. 77, 5705–5705, DOI: 10.1158/1538-7445.AM2017-5705 (2017).

16. Plagnol, V. et al. Analytical validation of a next generation sequencing liquid biopsy assay for high sensitivity broad molecular profiling. PLoS ONE 13, e0193802, DOI: 10.1371/journal.pone.0193802 (2018).

17. Kinde, I., Wu, J., Papadopoulos, N., Kinzler, K. W. & Vogelstein, B. Detection and quantification of rare mutations with massively parallel sequencing. Proc. Natl. Acad. Sci. 108, 9530–9535, DOI: 10.1073/pnas.1105422108 (2011). arXiv:1408.1149.

18. Shiroguchi, K., Jia, T. Z., Sims, P. A. & Xie, X. S. Digital RNA sequencing minimizes sequence-dependent bias and amplification noise with optimized single-molecule barcodes. Proc. Natl. Acad. Sci. 109, 1347–1352, DOI: 10.1073/pnas. 1118018109 (2012). arXiv:1408.1149.

19. Hindson, B. J. et al. High-throughput droplet digital PCR system for absolute quantitation of DNA copy number. *Anal*. chemistry 83, 8604–10, DOI: 10.1021/ac202028g (2011).

20. Kline, M. C., Romsos, E. L. & Duewer, D. L. Evaluating Digital PCR for the Quantification of Human Genomic DNA: Accessible Amplifiable Targets. Anal. Chem. 88, 2132–2139, DOI: 10.1021/acs.analchem.5b03692 (2016).

21. American College of Obstetricians and Gynecologists. ACOG committee opinion: Carrier Screening for Genetic Conditions. Tech. Rep. (2017).

22. Prior, T. W. Carrier screening for spinal muscular atrophy. Genet. Medicine 10, 840–842, DOI: 10.1097/GIM.0b013e318188d069 (2008).

23. Williams, T. N. & Weatherall, D. J. World distribution, population genetics, and health burden of the hemoglobinopathies. Cold Spring Harb. Perspectives Medicine 2, a011692, DOI: 10.1101/cshperspect.a011692 (2012).

24. https://www.cdc.gov/ncbddd/sicklecell/data.html.

25. Ojodu, J. Incidence of Sickle Cell Trait — United States, 2010. Centers for Dis. Control. Prev. MMWR 63, 1155–1158, DOI: 10.1242/jeb.02455 (2014).

26. Strom, C. M. et al. Cystic fibrosis testing 8 years on: Lessons learned from carrier screening and sequencing analysis. Genet. Medicine 13, 166–172, DOI: 10.1097/GIM.0b013e3181fa24c4 (2011).

27. Muralidharan, K. et al. Population carrier screening for spinal muscular atrophy: A position statement of the association for molecular pathology, DOI: 10.1016/j.jmoldx.2010.11.012 (2011).

28. Akolekar, R., Beta, J., Picciarelli, G., Ogilvie, C. & D’Antonio, F. Procedure-related risk of miscarriage following amniocentesis and chorionic villus sampling: A systematic review and meta-analysis. Ultrasound Obstet. Gynecol. 45, 16–26, DOI: 10.1002/uog.14636 (2015).

29. Snyder, M. W., Kircher, M., Hill, A. J., Daza, R. M. & Shendure, J. Cell-free DNA Comprises an in Vivo Nucleosome Footprint that Informs Its Tissues-Of-Origin. Cell 164, 57–68, DOI: 10.1016/j.cell.2015.11.050 (2016). 15334406.

30. Sinha, R. et al. Index Switching Causes “Spreading-Of-Signal” Among Multiplexed Samples In Illumina HiSeq 4000 DNA Sequencing. bioRxiv 125724, DOI: 10.1101/125724 (2017).

31. Vodák, D. et al. Sample-Index Misassignment Impacts Tumour Exome Sequencing. Sci. Reports 8, 5307, DOI: 10.1038/s41598-018-23563-4 (2018).

32. MacConaill, L. E. et al. Unique, dual-indexed sequencing adapters with UMIs effectively eliminate index cross-talk and significantly improve sensitivity of massively parallel sequencing. BMC Genomics 19, 30, DOI: 10.1186/s12864-017-4428-5 (2018).

33. Minniti, C. P. et al. Elevated tricuspid regurgitant jet velocity in children and adolescents with sickle cell disease: Association with hemolysis and hemoglobin oxygen desaturation. Haematologica 94, 340–347, DOI: 10.3324/haematol.13812 (2009).

34. Camunas-Soler, J. et al. Noninvasive prenatal diagnosis of single-gene disorders by use of droplet digital PCR. Clin. Chem. 64, 336–345, DOI: 10.1373/clinchem.2017.278101 (2018).

35. Lun, F. M. F. et al. Noninvasive prenatal diagnosis of monogenic diseases by digital size selection and relative mutation dosage on DNA in maternal plasma. Proc. Natl. Acad. Sci. 105, 19920–19925, DOI: 10.1073/pnas.0810373105 (2008). arXiv:1408.1149.

36. Mohanty, D. et al. Prevalence of *β*-thalassemia and other haemoglobinopathies in six cities in India: A multicentre study. J. Community Genet. 4, 33–42, DOI: 10.1007/s12687-012-0114-0 (2013).

37. Devonshire, A. S. et al. Towards standardisation of cell-free DNA measurement in plasma: Controls for extraction efficiency, fragment size bias and quantification. Anal. Bioanal. Chem. 406, 6499–6512, DOI: 10.1007/s00216-014-7835-3 (2014).

38. Haque, I. S. et al. Modeled fetal risk of genetic diseases identified by expanded carrier screening. JAMA - J. Am. Med. Assoc. 316, 734–742, DOI: 10.1001/jama.2016.11139 (2016).

39. Mincy, R., Garfinkel, I. & Nepomnyaschy, L. In-hospital paternity establishment and father involvement in fragile families. J. Marriage Fam. 67, 611–626, DOI: 10.1111/j.1741-3737.2005.00157.x (2005).

40. Bellis, M. A., Hughes, K., Hughes, S. & Ashton, J. R. Measuring paternal discrepancy and its public health consequences, DOI: 10.1136/jech.2005.036517 (2005).

41. Barrett, A. N., McDonnell, T. C., Chan, K. C. & Chitty, L. S. Digital PCR analysis of maternal plasma for noninvasive detection of sickle cell anemia. Clin. Chem. 58, 1026–1032, DOI: 10.1373/clinchem.2011.178939 (2012).

42. Lun, F. M. F. et al. Noninvasive prenatal diagnosis of monogenic diseases by digital size selection and relative mutation dosage on DNA in maternal plasma. Proc. Natl. Acad. Sci. 105, 19920–19925, DOI: 10.1073/pnas.0810373105 (2008). arXiv:1408.1149.

43. Kivioja, T. et al. Counting absolute numbers of molecules using unique molecular identifiers. Nat. Methods 9, 72–74, DOI: 10.1038/nmeth.1778 (2012).

44. Aigrain, L., Gu, Y. & Quail, M. A. Quantitation of next generation sequencing library preparation protocol efficiencies using droplet digital PCR assays - a systematic comparison of DNA library preparation kits for Illumina sequencing. BMC Genomics 17, 458, DOI: 10.1186/s12864-016-2757-4 (2016).

45. Kou, R. et al. Benefits and challenges with applying unique molecular identifiers in next generation sequencing to detect low frequency mutations. PLoS ONE 11, e0146638, DOI: 10.1371/journal.pone.0146638 (2016).

46. Zarrei, M., MacDonald, J. R., Merico, D. & Scherer, S. W. A copy number variation map of the human genome, DOI: 10.1038/nrg3871 (2015).

47. Beroukhim, R. et al. The landscape of somatic copy-number alteration across human cancers. Nature 463, 899–905, DOI: 10.1038/nature08822 (2010).

48. Van der Auwera, G. A. et al. From fastQ data to high-confidence variant calls: The genome analysis toolkit best practices pipeline. Curr. Protoc. Bioinforma. 43, 11.10.1–11.10.33, DOI: 10.1002/0471250953.bi1110s43 (2013). NIHMS150003.

